# Circulating proteomic patterns in AF related left atrial remodeling indicate involvement of coagulation and complement cascade

**DOI:** 10.1101/327759

**Authors:** Jelena Kornej, Petra Büttner, Elke Hammer, Beatrice Engelmann, Borislav Dinov, Philipp Sommer, Daniela Husser, Gerhard Hindricks, Uwe Völker, Andreas Bollmann

## Abstract

**Background:** Left atrial (LA) electro-anatomical remodeling and diameter increase in atrial fibrillation (AF) indicates disease progression and is associated with poor therapeutic success. Furthermore, AF leads to a hypercoagulable state, which in turn promotes the development of a substrate for AF and disease progression in the experimental setting. The aim of this study was to identify pathways associated with LA remodeling in AF patients using untargeted proteomics approach.

**Methods:** Peripheral blood samples of 48 patients (62±10 years, 63% males, 59% persistent AF) undergoing AF catheter ablation were collected before ablation. 24 patients with left atrial low voltage areas (LVA), defined as <0.5 mV, and 24 patients without LVA were matched for age, gender and CHA_2_DS_2_-VASc score. Untargeted proteome analysis was performed using LC-ESI-Tandem mass spectrometry in a label free intensity based workflow. Significantly different abundant proteins were identified and used for pathway analysis and protein-protein interaction analysis.

**Results:** Analysis covered 280 non-redundant circulating plasma proteins. The presence of LVA correlated with 30 differentially abundant proteins of coagulation and complement cascade (q<0.05).

**Conclusions:** This pilot proteomic study identified plasma protein candidates associated with electro-anatomical remodeling in AF and pointed towards an imbalance in coagulation and complement pathway, tissue remodeling and inflammation

## Introduction

Atrial fibrillation (AF) is the most common sustained arrhythmia occurring in approximately 2% of the general population [1]. It is a progressive disease as an evolution from paroxysmal to persistent AF (AF type) is frequently observed. This is related to advanced structural and electrical left atrial remodeling. Structural remodeling might be suggested by electrocardiography as increased left atrial diameter (LAD). Evidence for electro-anatomical remodeling is currently detectable only invasively during catheter ablation in up to 20–25% of AF patients [2] and is represented by low voltage areas (LVA), e.g. pathologically low electrical signals with amplitudes less than 0.5 mV. Previous studies suggested that LVA correspond to areas of fibrotic and electrically silent myocardium [3]. Advanced LA remodeling is associated with worse rhythm outcomes after catheter ablation [2,4,5] and as consequence is associated with higher rates of repeated catheter ablations.

One of the most serious complications in AF patients related to hypercoagulability and impaired blood flow is an ischemic stroke [6,7] also leading to increased hospitalization and cost-intensive treatment.^1^ The increased risk for thromboembolic events is attributable to alterations in blood flow combined with anatomical and structural defects resulting in the fulfilment of Virchow’s triad for thrombogenesis and a hypercoagulable state in AF patients [8]. Interestingly, recent experimental studies indicated that hypercoagulability promotes the development of an electro-anatomical substrate and AF progression [9].

Because LVA detection is applicable as standardized procedure only during AF catheter ablation, this led to an increased research interest in biomarker profiles that mirror AF progression and can be used to predict rhythm outcome and AF-related complications non-invasively [10–12]. In the current study we applied a novel approach, where we performed an untargeted proteome screening of high and medium abundant circulating blood proteins to identify differentially expressed proteins in AF patients with and without LVA and with LAD ≥ or < 44mm. Furthermore, the proteins were used to characterize potential pathophysiological pathways associated with LA remodeling in AF patients.

## Methods

### Study population

The study population comprised 48 patients with symptomatic AF who underwent catheter ablation at Heart Center Leipzig, Germany. Twenty-four patients with LVA and without LVA respectively were matched according to age, gender and CHA_2_DS_2_-VASc score. The study was approved by the local Ethical Committee (Medical Faculty, University Leipzig) and patients provided written informed consent for participation. Paroxysmal and persistent AF were defined according to current guidelines [13]. Paroxysmal AF self-terminated within 7 days after onset. Persistent AF lasted longer than 7 days or required drugs or direct current cardioversion for termination. In all patients, transthoracic and transesophageal echocardiography were performed prior to the ablation. LAD was measured in parasternal long axis view in M-Mode. All class I or III antiarrhythmic medications with exception of amiodarone were discontinued for at least 5 half-lives before the AF ablation.

### Electro-anatomical mapping and definition of substrate

The electro-anatomical mapping was performed in sinus rhythm. In patients presenting with atrial fibrillation, the arrhythmia was terminated by electrical cardioversion and the mapping was further performed in sinus rhythm. End-point of the catheter ablation was isolation of the PV with proof of both exit and entrance block. The electro-anatomical voltage maps of the LA excluding the pulmonary veins were created using multielectrode spiral catheter with interelectrode distance 2-5-2 or ablation catheter with a 3.5 mm electrode tip and contact measurement properties (SmartTouch Thermocool and TactiCath, St Jude Medical (SJM), Saint Paul, MN, USA) as mapping catheter. Electro-anatomical mapping was performed using 3-D electro-anatomical mapping systems (Carto, Biosence Webster, Diamond Bar, CA, USA or EnSite Precision, SJM). Electrograms with amplitudes over 0.5 mV were defined as normal potentials, and signals with amplitude under 0.5 mV as low-voltage potentials. In areas with low-voltage amplitudes we aimed at sufficient mapping points density (>200 points). Points with insufficient catheter-to-tissue contact or inside ablation lines were excluded.

At the end of the procedure, an attempt to induce AF or left atrial macro-reentry tachycardia (LAMRT) was performed using a standardized protocol (burst stimulation with 300, 250, 200 ms from coronary sinus). According to the underlying LVA and inducible LAMRT additional ablation lines were applied.

### Blood samples

Blood samples were obtained in EDTA test tubes in fasting state prior ablation procedure from femoral vein and processed within one hour after collection. Blood plasma was prepared (1000 × g for 10 min at 20°C) and aliquots were stored at −70 °C for subsequent analyses.

### Proteomic analysis

Samples were prepared, measured by mass spectrometry and data analyzed according to a modified protocol described earlier [14]. Briefly, plasma samples were subjected to immunoaffinity subtraction of six high abundant proteins using the Multiple Affinity Removal System (MARS6) (Agilent Technologies, Wilmington, DE, USA). Protein in the flow through fraction was precipitated using trichloroacetic acid (TCA) at a final concentration of 15% (v/v). Precipitates were reconstituted in urea/thiourea (8M/2M) and protein was determined using a Bradford assay (Bio-Rad, Hercules, CA, USA). Four µg protein lysate were reduced and alkylated and digested with trypsin (Promega, Madison, WI, USA) with a ratio of 1:25 at 37°C overnight. Peptide purification was performed with μC18 ZipTip^®^ (Millipore Cooperation, Billerica, MA, USA) as described before [15]. LC-ESI tandem mass spectrometry was carried out on a nanoAquity UPLC (Waters) - LTQ Orbitrap Velos MS (Thermo Scientific Electron, Bremen, Germany) configuration. High precision precursor scans were recorded in the Orbitrap and MS/MS data were recorded in data-dependent mode for the Top-20 peaks in the ion trap.

Raw MS data were processed using the Refiner MS 10.0 and Analyst 10.0 module (Genedata, Basel, Switzerland). Identification was performed against a human FASTA-formatted database containing 20,155 unique sequence entries (reviewed human database, release of 2016/06) with MASCOT (v2.3.2, Matrix Science, London, UK) as search engine. Carbamidomethylation of cysteine was set as fixed modification and oxidation of methionine as variable modification. Enzyme specificity was selected to trypsin with using 10 ppm MS tolerance and 0.6 Da MS/MS tolerances. Peptide FDR was set to 1% of protein groups identified. Peptides sharing protein group identifications within the data set were not considered for quantitative analyses.

Relative quantification protein values were exported from Refiner as summed intensities of Hi3 peptides per protein. Preprocessing steps before statistical analysis included removal of proteins with more than 50 % missing values per group, which diminished the initially identified 280 to 237 proteins, and log10 transformation of data.

All methods were performed in accordance with the relevant guidelines and regulations.

### Statistical analysis

Data are presented as mean and standard deviation (SD) if normally distributed or as median [interquartile range] for skewed continuous variables and as proportions for categorical variables. Continuous variables were tested for normal distribution using the Kolmogorov-Smirnov test. The differences between continuous values were assessed using an unpaired t-test or the Mann-Whitney, and a chi-square test for categorical variables. A p-value <0.05 was considered statistically significant, and all analyses were performed with SPSS statistical software version 23. Proteomic data significance tests were corrected for multiple testing using Benjamini-Hochberg false discovery rate and significance cut-off set to *q*<0.05.

We detected differentially expressed proteins by comparing patients with LVA vs. without LVA (24 patients in each group) and patients with LAD<44mm (n=20) and LAD≥44mm (n=28).

Categorization of proteins in particular physiological pathways was done using WebGestalt [16] accessing databases of Kyoto Encyclopedia of Genes and Genomes (KEGG) [17] and gene ontology (GO) [18]. Over-enrichment p-values are not reported as the applied screening method only detects highly abundant plasma proteins and an unbiased over-enrichment analysis of the plasma wide proteome is not possible.

## Results

Untargeted proteomic analysis was performed in 48 AF patients matched for age, gender and CHA_2_DS_2_-VASc score). The clinical characteristics of study population are presented in Table 1. There were no differences in age, gender, AF type, BMI, renal function or CHA_2_DS_2_-VASc score. However, patients with LVA had significantly larger LAD than patients without (p=0.01).

**Table 1.** Clinical characteristics of study population

|  | Low voltage areas |  | p-value |
| --- | --- | --- | --- |
|  | No (n=24) | Yes (n=24) |  |
| Age, y | 66 ± 10 | 67 ± 11 | 0.074 |
| Males, % | 64 | 65 | 0.930 |
| Persistent AF, % | 68 | 74 | 0.653 |
| BMI, kg/m<sup>2</sup> | 30 ± 5 | 30 ± 4 | 0.657 |
| eGFR, ml/min/1.73m<sup>2</sup> | 72 ± 14 | 66 ± 20 | 0.218 |
| CHA<sub>2</sub>DS<sub>2</sub>-VASc score | 2.8 ± 1.3 | 3.3 ± 1.6 | 0.240 |
| LA Diameter, mm | 42 ± 6 | 47 ± 7 | 0.010 |
| EF, % | 56 ± 7 | 52 ± 12 | 0.201 |

### Proteomic analysis

Analysis covered 1.985 unique peptides representing 280 non-redundant proteins (see supplementary data table 1). Among the 237 proteins being quantified in at least 50 % of the patients in each group, 30 proteins displayed different levels (t-test (FDR), p<0.05) in patients presenting with LVA in comparison to the control group (Table 2). L-selectin and peptidase inhibitor 16 had higher abundances in the control group with fold changes > 1.5. Ten proteins were identified to be >1.5-fold more abundant in the LVA group. Tetratricopeptide repeat protein 39A and SWI/SNF complex subunit SMARCC1 were more than 2-fold enriched in plasma from this group.

**Table 2:**
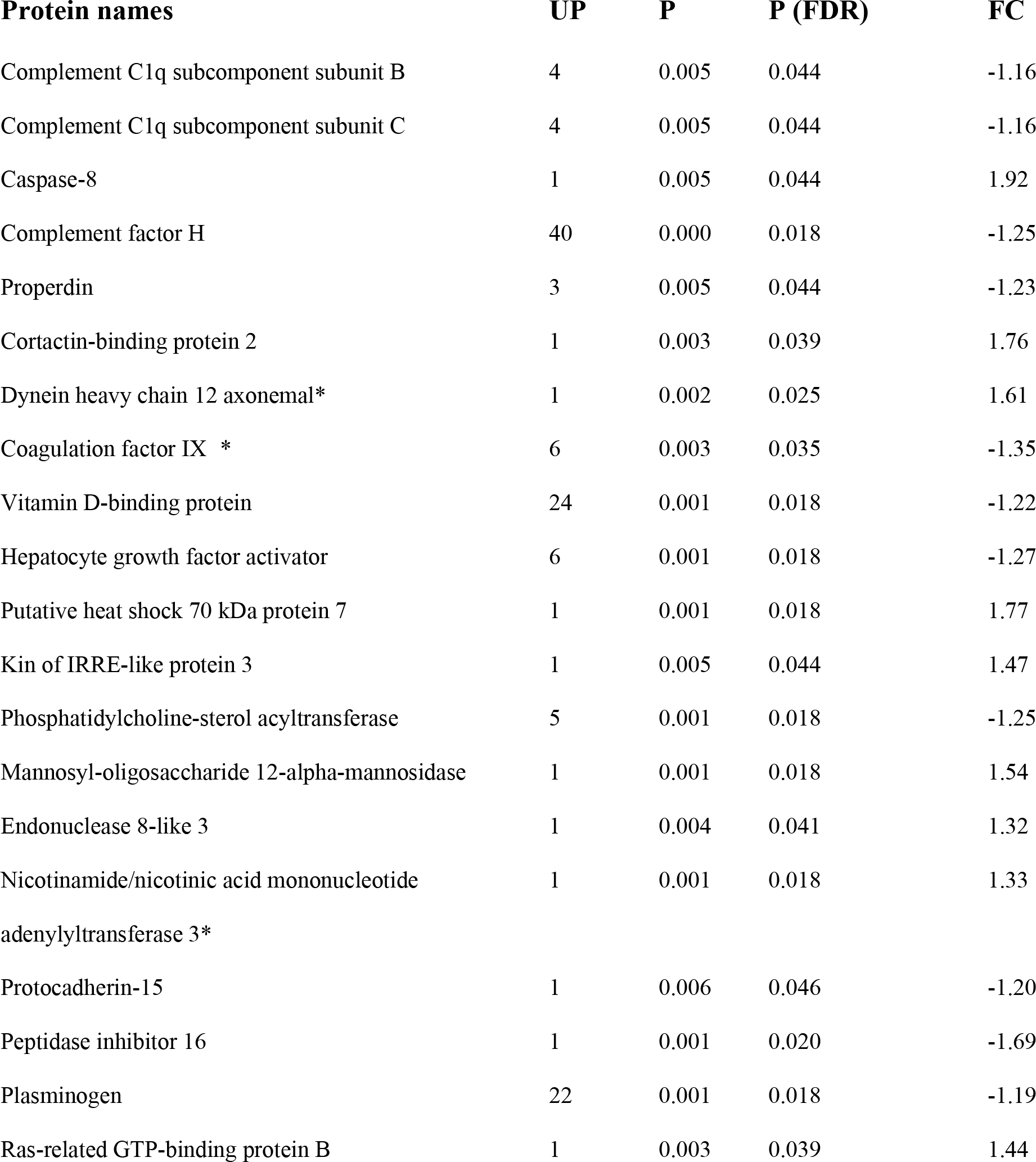
Significantly differentially expressed circulating plasma proteins in AF patients with-compared to patients without LVA. UP - unique peptides of the protein identified in proteome screening, P - t-test p-value, P(FDR) - p-value with FDR, FC - fold change with LVA vs. without LVA, negative fold change indicates lower concentrations in the LVA group. An asterisk marks proteins that were also differentially abundant in patients with LAD<44mm vs. LAD≥44mm (significance in t-test but not following FDR).

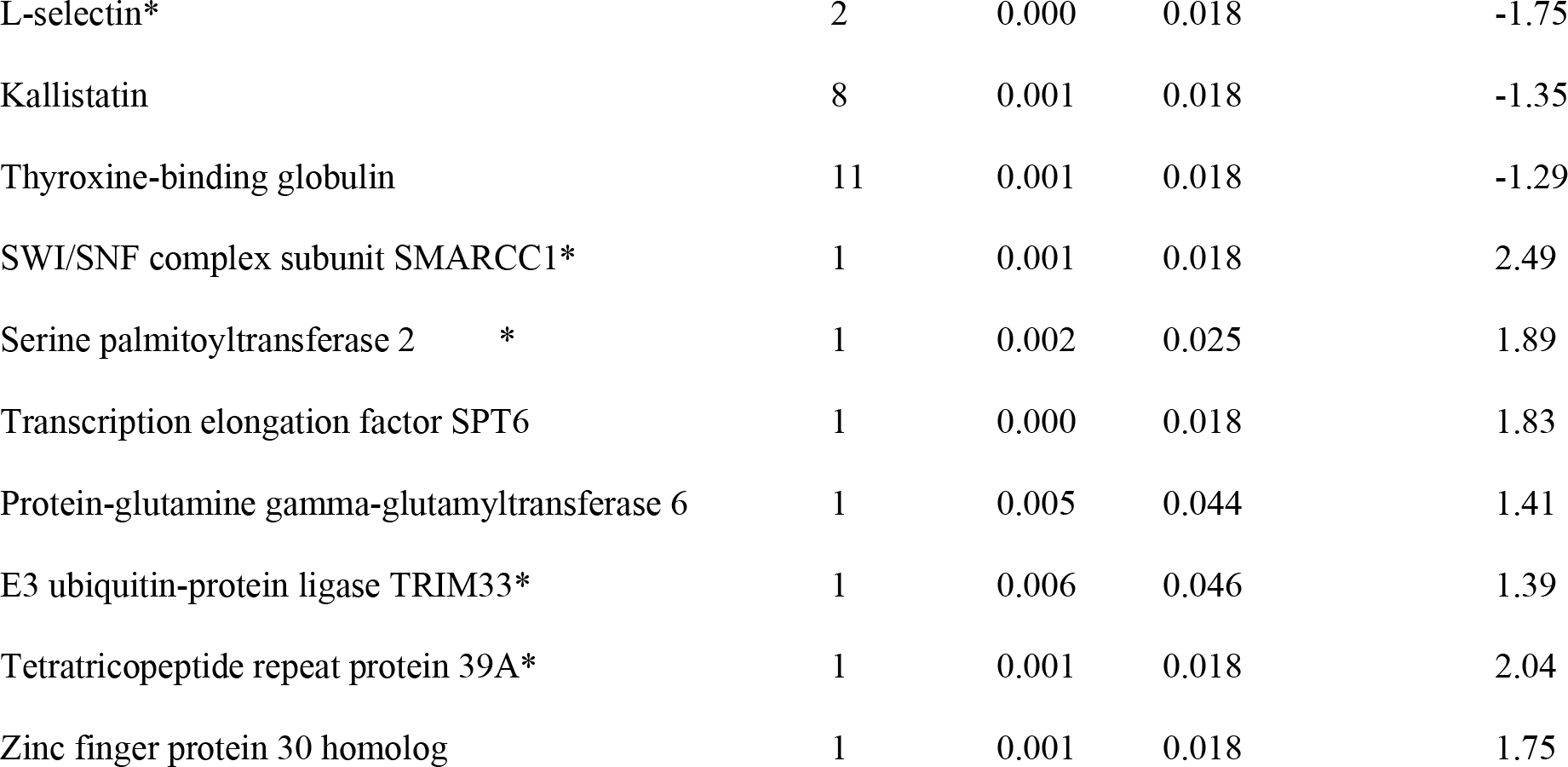

We annotated all 30 candidate proteins to physiological pathways using WebGestalt [16] to get an insight into physiological impact. F9, CFH, CFP, C1QB, and C1Q were assigned to GO:0072376 Protein activation cascade. CASP8, PI16, SERPINA4, SMARCC1, and SERPINA7 were assigned to GO:0045861 Negative regulation of proteolysis. CFH, CFP, C1QB, and C1QC were assigned to GO:0006959 Humoral immune response. F9, CFH, PLG, C1QB, and C1QC were assigned to KEGG:hsa04610 Complement and coagulation cascades. When comparing LAD<44mm and LAD>44mm 26 proteins were found differentially expressed in t-test but did not pass multiple testing corrections (Table 2 and supplement Table 2).

## Discussion

This study used an untargeted proteomic approach to discover high and medium abundant proteins related to left-atrial remodeling in AF patients. Significant differences in protein profiles in patients with and without LVA were detected. The pathway analysis suggests that alterations in the complement and coagulation cascade, inflammatory processes, and tissue remodeling are associated with LVA and LAD. We therefore assume that the observed alterations are more specific for LVA than for increased left diameter although both phenotypes are interrelated. This is in line with previous findings [4] and of great clinical importance. However, because of relatively small cohort, proteomic differences in LAD phenotype did not reach significance after FDR

### Complement and coagulation cascade and electro-anatomical remodeling

As initially stated, hypercoagulability associated with an increased risk for stroke, is a common finding in AF patients [6]. In our study, we observed differential abundance of proteins from the complement and coagulation system. Classical complement cascade is started by activation of multimolecular C1-complex which consists of C1q, C1r, and C1s. We found C1q-B and C1q-C significantly lower in the patients with LVA, whereas the third C1q subcomplex C1q-A was detected but unchanged. C1qr as well as C1qs were also found to be lower in LVA patient plasma but did not pass FDR significance testing (see supplement Table 1). Decreased C1-complex activity was found to associate with impaired immune response and auto-immune reactions [19]. CFH and properdin were also less abundant in the patients with LVA. Both are complement system regulators which identify and protect the host from complement also during pathological states. An imbalance might therefore contribute to tissue damage [20,21].

Plasminogen typically is highly abundant in blood and if catalyzed into plasmin is fibrinolytic. Downregulation of plasminogen, which might explain the lower levels observed in LVA patients, could therefore attribute to the hypercoagulation state often observed in AF. F9 is part of the intrinsic coagulation cascade, is activated through factor 11, and then cleaves factor 10 which facilitates prothrombin cleavage. Prothrombin was found to be lower in LVA patient plasma but did not pass FDR significance testing (see supplement Table 1). Altogether, our data indicate an imbalance of coagulation and complement cascade activation, homeostasis, and regulation in AF patients with LVA and generate hypothesis regarding whether progression of electro-anatomical remodeling is cause or consequence to those alterations.

### Fibrosis, inflammation, and electro-anatomical remodeling

LVA is supposed to be an electro-anatomical remodeling with fibrotic character. Fibrosis is characterized by increased production of ECM, cell death, cell trans-differentiation e.g. myofibroblast activation and infiltration of inflammatory cells [22]. Several proteins found altered by our approach are known to contribute to those pathological processes. The highest plasma level decrease in the LVA group was found for cell adhesion molecule L-selectin. L-selectin is expressed on a variety of leukocytes, but is best explored in naïve T-lymphocytes where its shedding from the cell membrane in the primary lymph nodes occurs during their activation and has implications for subsequent cell fate. Shedding from the surface is thought to mobilize intracellular cytotoxic granules leading to the release of lytic enzymes towards target cells [23]. Interestingly, L-selectin was also proposed to be a receptor for the earlier discussed CFH, thus connecting immune function and complement system [24]. Furthermore, L-selectin was found to be down regulated through shedding by atrial natriuretic peptide (ANP) leading to decreased neutrophil infiltration [25]. Recently we found that plasma ANP levels are significantly higher and therewith predictive for LVA in AF patients (data in submission) what is well in line with the literature reports and the now observed lower levels of L-selectin.

Furthermore, we observed altered levels of proteins that are annotated to G0:0045861 Negative regulation of proteolysis whereas proteolysis is indispensable in fibrosis and inflammation [26]. Peptidase inhibitor 16 (PI16), which was lower in the LVA group, was assigned to this group. PI16 might contribute to electro-anatomical remodeling as the protein is known to be regulated by sheer stress, to inhibit ECM-remodeling matrix metalloproteinases, and is thought to maintain anti-inflammatory and non-activated cell homeostasis [27]. In context of heart failure, PI16 is supposed to inhibit myocyte hypertrophy [27]. Caspase-8, another proteolytic enzyme, is involved in controlled extrinsic apoptosis induced via death receptor. The observed increase in the LVA patient plasma might represent apoptotic activity. Higher circulating levels of caspase-8 were found to associate with increased incidence of cardiovascular events [28]. Several proteins found at higher abundance in LVA patients are involved in transcriptional processes or chromatin remodeling respectively, namely SWI/SNF complex subunit SMARCC1 (SMARCC1), transcription elongation factor SPT6 (SUPT6H), and zinc finger protein 30 homologue (ZFP30). Chromatin remodeling was already identified as promising tool for intervention in fibrotic and inflammatory disease (YANG, targeting). SMARCC1 showed the highest observed difference between the experimental groups in our screening. SMARCC1 is component of a chromatin remodeling complex which is needed to induce transcription e.g. in neurogenesis [29]. ZFP30 encodes a transcription factor which was found to be involved in differentiation processes and inflammatory response [30]. SUPT6H alters nuclear histone status to allow for a transcriptionally competent chromatin state [31] especially important for expression of genes in developmental and differentiation e.g. myogenesis [32].

The second most altered protein in our study was tetratricopeptide repeat protein 39A (TTC39A) whose function is widely unknown. Interestingly, an interaction of TTC39A with P2RY12, a receptor involved in platelet aggregation, regarded to be a potential target of thromboembolism treatment, was supposed [33].

Altogether, we suppose that the observed markers associate with altered cell proliferation rates or cell differentiation processes.

## Limitations

Several proteins that were discussed typically act inside cells or are bound to them. We interpret this to be indicative for shedding, secretion, endothelial damage (e.g. in consequence of increased atrial sheer stress) or cell death (e.g. fibrosis-related decline of cardiomyocytes). Although, sample processing aimed to limit cell contamination of the plasma used in this study the contamination with cell debris is unlikely, nevertheless, it cannot be excluded.

The used technology only detected proteins with high and medium abundance in plasma; typically, 250–350 proteins with concentrations in a range of μg to mg per ml. Lower concentrations were not detected whereas we cannot estimate the quantity of natriuretic peptides, growth factors or cytokines. This limitation also impedes over-enrichment analysis of physiological pathways. We used pathway annotation tools to get insights into the physiological pathways the identified candidates are assigned to.

In general, the source of the detected proteins is beyond the scope of this study and causal association with atrial LVA cannot be proved. Further larger studies are needed to address this aspect.

## Conclusions

This pilot proteomic study identified plasma protein candidates associated with electro-anatomical remodeling in AF and pointed towards an involvement of coagulation and complement pathway, inflammation, and tissue remodeling. Further studies are underway to replicate and apply these findings.

## Supporting information

**S1 Table 1**: Proteins detected in proteomic approach comparing 24 patients with LVA and 24 patients without LVA. UP - number of unique peptides belonging to the protein which were detected in the screening, p-value - from T-Test comparing LVA and controls, p-value (FDR) - Benjamini Hochberg false discovery rate, FC - fold change in protein levels. Proteins in bold were found significantly different in LVA group compared to controls.

